# Spectral Jaccard Similarity: A new approach to estimating pairwise sequence alignments

**DOI:** 10.1101/800581

**Authors:** Tavor Z. Baharav, Govinda M. Kamath, David N. Tse, Ilan Shomorony

## Abstract

A key step in many genomic analysis pipelines is the identification of regions of similarity between pairs of DNA sequencing reads. This task, known as pairwise sequence alignment, is a heavy computational burden, particularly in the context of third-generation long-read sequencing technologies, which produce noisy reads. This issue is commonly addressed via a two-step approach: first, we filter pairs of reads which are likely to have a large alignment, and then we perform computationally intensive alignment algorithms only on the selected pairs. The *Jaccard similarity* between the set of *k*-mers of each read can be shown to be a proxy for the alignment size, and is usually used as the filter. This strategy has the added benefit that the Jaccard similarities don’t need to be computed exactly, and can instead be efficiently estimated through the use of *min-hashes*. This is done by hashing all *k*-mers of a read and computing the minimum hash value (the min-hash) for each read. For a randomly chosen hash function, the probability that the min-hashes are the same for two distinct reads is precisely their *k*-mer Jaccard similarity. Hence, one can estimate the Jaccard similarity by computing the fraction of min-hash collisions out of the set of hash functions considered.

However, when the *k*-mer distribution of the reads being considered is significantly non-uniform, Jaccard similarity is no longer a good proxy for the alignment size. In particular, genome-wide GC biases and the presence of common *k*-mers increase the probability of a min-hash collision, thus biasing the estimate of alignment size provided by the Jaccard similarity. In this work, we introduce a min-hash-based approach for estimating alignment sizes called *Spectral Jaccard Similarity* which naturally accounts for an uneven *k*-mer distribution in the reads being compared. The Spectral Jaccard Similarity is computed by considering a min-hash collision matrix (where rows correspond to pairs of reads and columns correspond to different hash functions), removing an offset, and performing a *singular value decomposition*. The leading left singular vector provides the Spectral Jaccard Similarity for each pair of reads. In addition, we develop an approximation to the Spectral Jaccard Similarity that can be computed with a single matrix-vector product, instead of a full singular value decomposition.

We demonstrate improvements in AUC of the Spectral Jaccard Similarity based filters over Jaccard Similarity based filters on 40 datasets of PacBio reads from the NCTC collection. The code is available at https://github.com/TavorB/spectral_jaccard_similarity.

## 1 Introduction

The advent of long-read sequencers such as PacBio and Oxford Nanopore have made the goal of obtaining gold-standard genome assemblies a reality. Unlike short read technologies, which provide reads of length 100-200 bp with an error rate of 1%, chiefly substitution errors, long read technologies provide reads of lengths in the tens of thousands with a nominal error rate of 13-15%, consisting mostly of insertions and deletions [Weirather et al., 2017]. While the long reads make resolving repeated sequences easier, the higher error rates make the computational tasks required for assembly significantly more challenging.

Genome assembly is usually performed based on one of two main approaches: *de-novo* assembly, where one attempts to assemble a new genome “from scratch” using only the reads obtained, and *reference-based* assembly, where one assembles the reads using a pre-assembled genome of a related organism. Alignment is an integral part of most pipelines in either approach, and is often the most time consuming step (see Fig. S18). In both settings naive dynamic programming based alignment [Needleman and Wunsch, 1970, Smith and Waterman, 1981, Myers, 1986] is impractical due to its quadratic time complexity.

In reference-based assembly pipelines [Vaser et al., 2017, Wick, 2019], where one has a reference of a related organism, the first step usually consists of aligning all reads to the reference; i.e., read-to-reference alignment. For *n* reads each of length *L* and a reference of length *G*, the time complexity of aligning all reads to the reference via dynamic programming is *O*(*nLG*), which is impractical in settings where there are *n* ~ 10^5^ reads, each of length *L* ~ 10^4^, with *G* on the order of 10^6^-10^11^ depending on the organism.

Similarly, the first step in most de-novo assembly pipelines [Chin et al., 2013, Berlin et al., 2015, Li, 2016, Chin et al., 2016, Koren et al., 2017, Kamath et al., 2017] is the pairwise alignment of all reads, which is computationally very costly. For *n* reads each of length *L*, the time complexity of aligning all pairs of reads would be *O*(*n*^2^*L*^2^). Even for bacterial genome datasets where the number of reads obtained is on the order of *n* ~ 10^5^, with *L* ~ 10^4^, this is impractical. For most of this work we discuss alignment in the context of pairwise read alignment. However, our ideas can be adapted to the read-to-reference alignment paradigm.

One key observation that helps alleviate this computational burden is that in practice one only cares about alignment between reads when there is a significant overlap. Further, as shown in Figure 4(a), in a typical dataset, more than 99.99% of pairs of reads do not have a significant overlap. Hence, most practical read aligners follow a *seed-and-extend* paradigm. The “seeding” step typically involves identifying pairs of reads that share many *k*-mers (length-*k* contiguous substrings). This step can be understood as a way to “filter” the set of read pairs in order to select those that share a reasonable number of *k*-mers and are thus likely to have a significant overlap [Myers, 2014, Berlin et al., 2015, Li, 2016, 2018]. Once these “candidate pairs” (whose number can be orders of magnitude smaller than the total number of read pairs) are obtained, computationally expensive dynamic-programming-based algorithms are used to obtain detailed alignment maps.

The idea of using the number of shared *k*-mers as a metric for filtering pairs of reads is equivalent to viewing the *Jaccard similarity* between the set of *k*-mers of each read as a proxy for the alignment size. Under standard implementations where computing set unions and set intersections has a linear time complexity in the sizes of the sets, this filtering step has a time complexity of *O*(*n*^2^*L*) for pairwise read alignment. Recently Jaccard similarity has been used in a variety of applications such as genome skimming [Denver et al., 2016], and in new methods to compare whole genomes and study taxonomic diversity in the microbiome [Ondov et al., 2016, Sarmashghi et al., 2019].

One very attractive property of Jaccard Similarity in the context of filtering pairs of reads is that this metric is amenable to efficient estimation through the use of *min-hashes*. This is done by hashing all *k*-mers on a read (the total number of *k*-mers in a length-*L* read is *L − k* + 1) and computing the minimum hash value (the min-hash) for each read. For a randomly chosen hash function, the collision probability for the min-hashes of two reads is precisely their Jaccard similarity. Hence, one can estimate the Jaccard similarity by computing the fraction of min-hash collisions out of the set of hash functions considered. For pairwise read alignment, if one uses *H* hashes to estimate the Jaccard similarity, this requires *O*(*nLH*) to compute the min-hashes and *O*(*n*^2^*H*) to compute the collisions giving us a time complexity of *O*(*n*^2^*H*) for the filtering step as generally, for regimes of interest, *n* ⨠ *L*. We note that this general approach is related to the minimizers method, which has been used both in the context of document fingerprinting and reducing storage requirements for biological sequence comparison [Schleimer et al., 2003, Roberts et al., 2004, Zheng et al., 2020] and to locality-sensitive hashing [Buhler, 2001, Marçais et al., 2019].

The idea of using min-hashes to estimate the Jaccard Similarity provides significant computational savings and is particularly effective when the genome where the reads come from is close to a random genome; i.e., a genome where every *k*-mer is equally likely to appear on a read. However, when the *k*-mer distribution of the reads being considered is significantly non-uniform, the Jaccard similarity is no longer a good proxy for the alignment size. In particular, genome-wide GC biases and the presence of common *k*-mers increase the probability of a min-hash collision, thus biasing the estimate of alignment size provided by the Jaccard similarity. In this work, we introduce a min-hash-based approach for estimating alignment sizes called *Spectral Jaccard Similarity*, which naturally accounts for an uneven *k*-mer distribution in the reads being compared. The Spectral Jaccard Similarity is computed by considering a min-hash collision matrix (where rows correspond to pairs of reads and columns correspond to different hash functions), removing an offset and performing a *singular value decomposition* (SVD). As illustrated in Figure 1, the leading left and right singular vectors can be seen as providing a rank-one approximation to the min-hash collision matrix. The leading left singular vector provides the Spectral Jaccard Similarity for each pair of reads, while the corresponding right singular vector can be understood as a measure of the “unreliability” or noise level of each hash function. Intuitively, a hash function that assigns low values to common *k*-mers is more unreliable for estimating alignment sizes, since it is more likely to create spurious min-hash collisions. Implicitly, this approach leads to a kind of weighted Jaccard similarity, where the weight of different hash functions is learned from the dataset.

**Figure 1:**
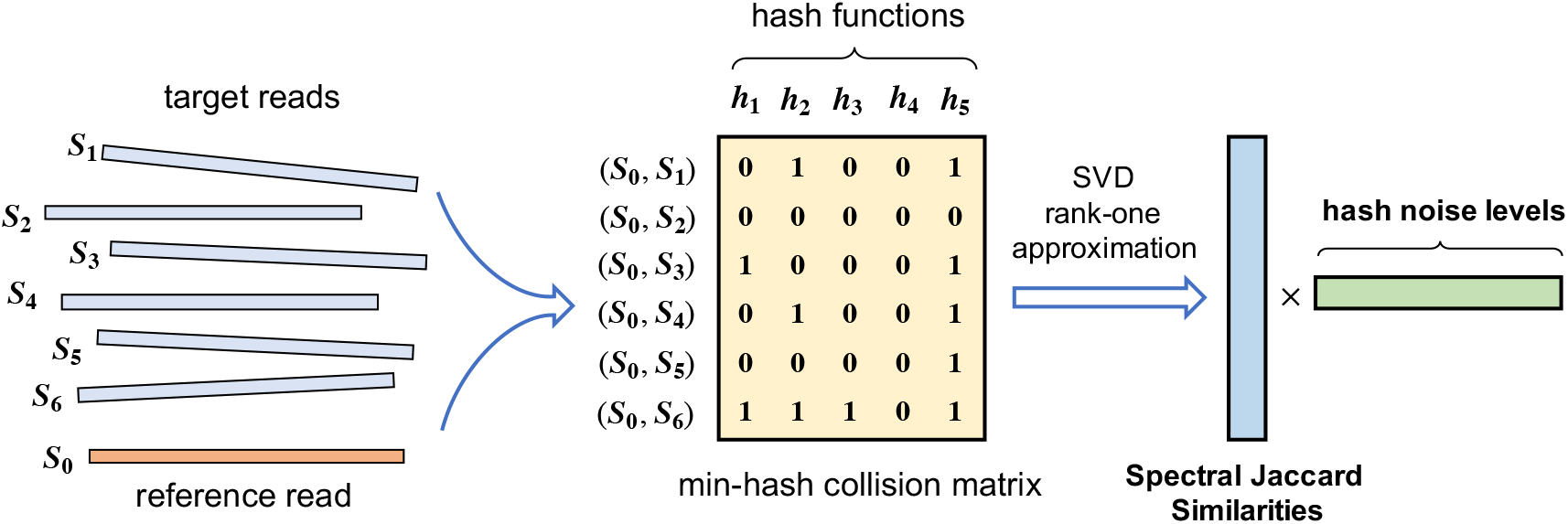
Overview of the Spectral Jaccard Similarity computation.

Experiments on PacBio long-read sequencing data from several bacterial genomes, spanning a variety of *k*-mer distributions, show that the Spectral Jaccard Similarity is significantly more correlated with alignment size than the standard Jaccard Similarity. When used as a metric to filter out pairs of reads that are unlikely to have a large alignment, Spectral Jaccard Similarity outperforms Jaccard Similarity on standard classification performance metrics. As an example, when applied to filtering pairs of reads which have an overlap of at least 30%, the area under the ROC curve (AUC) obtained by Spectral Jaccard Similarity filtering was consistently higher than the AUC obtained for Jaccard similarity on 40 datasets of the NCTC collection of Parkhill et al, as shown in Figure 2. These results are obtained using *k* = 7, which is an appropriate choice for the PacBio error rates [Myers, 2014].

**Figure 2:**
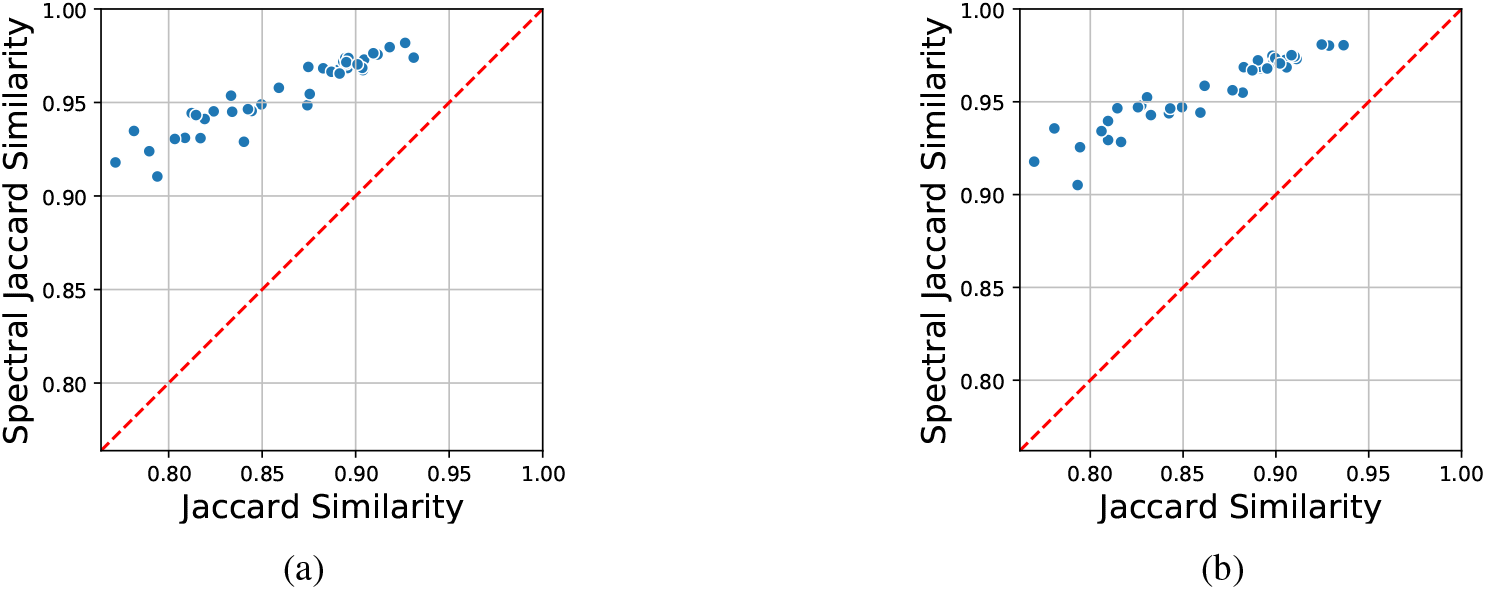
SJS has uniformly higher area under the ROC curve for experiments on 40 Pacbio bacterial datasets from the NCTC library [Parkhill et al]. In these experiments, Spectral Jaccard Similarity and Jaccard Similarity were used to filter pairs of reads with an overlap of at least 30%. SJS used was with 1000 hash functions, while Jaccard similarity was computed exactly. (*a*) shows the AUC values using Daligner alignments as ground truth. (*b*) shows the same results using Minimap2 alignments as ground truth.

This manuscript is organized as follows: in Section 2, we present a brief review of Jaccard similarity and its application to seed-and-extend algorithms for pairwise read alignment; in Section 3 we present the basis for Spectral Jaccard similarity and in Section 4 we show results on real and simulated datasets. We then conclude with a discussion in Section 5.

## 2 Jaccard Similarity

In general terms, the Jaccard similarity (JS) is a similarity metric between sets. For two sets *A* and *B*, the Jaccard similarity between them, JS(*A, B*), is defined as the size of their intersection divided by the size of their union. This is a very convenient measure as it is bounded between 0 and 1, JS(*A, B*) = 0 if and only if *A*⋂*B* = ∅, and JS(*A, B*) = 1 if and only if *A* = *B*. It has gained recent interest in its applications to finding documents (or web-pages) that are very similar but not the same as each other and in plagiarism detection. We refer the interested reader to Leskovec et al. [2014, Chapter 3] for a detailed review of the topic.

The Jaccard Similarity was applied to the problem of pairwise read alignment in Berlin et al. [2015], by considering the sets of *k*-mers of each read. For a fixed parameter *k*, the *k*-mer Jaccard similarity between reads *S*_0_ and *S*_1_ is given by

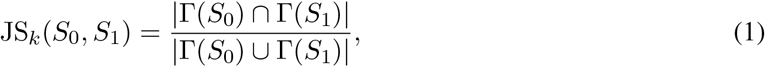

where Γ(*S*_*i*_) is defined as the set of *k*-mers for read *S*_*i*_. This is the same as *k*-shingle Jaccard similarity in the data mining literature [Manber et al., 1994, Broder, 1997]. In this case, JS can be viewed as a proxy for the size of the overlap (if any) between reads *S*_0_ and *S*_1_.

For instance, consider length-*L* reads *S*_0_ and *S*_1_ with an overlap of size *α*_0,1_*L*, for 0 ≤ *α*_0,1_ ≤ 1, as illustrated in Fig. 3, and let *p*_0,1_ be the fraction of overlap; i.e.,

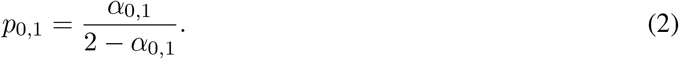

**Figure 3:**
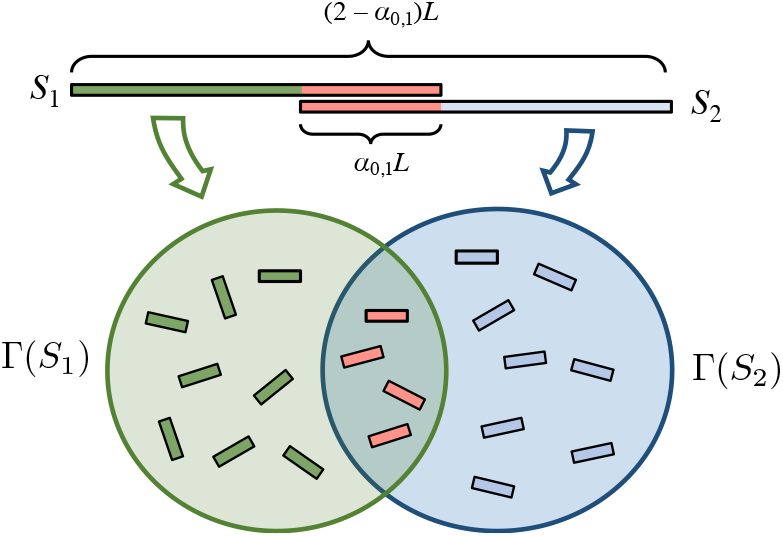
The *k*-mer Jaccard Similarity can be seen as a proxy for the alignment size.

If not many *k*-mers are shared by the non-overlapping parts of *S*_0_ and *S*_1_, we have that JS_*k*_(*S*_0_, *S*_1_) ≈ *p*_0,1_ making this a useful metric to filter pairs of reads with a high overlap. Notice that this approach is in a sense robust to errors in the reads. If we assume that each base is independently corrupted by noise (substitution, insertion or deletion) with probability *z*, then a *k*-mer is not corrupted with probability (1−*z*)^*k*^. Ignoring the unlikely event of collision of an erroneous *k*-mer with some other *k*-mer, JS_*k*_(*S*_0_, *S*_1_) is approximated as

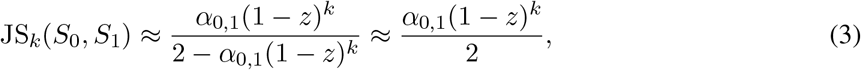

where the last approximation holds when (1−*z*)^*k*^ is small. Therefore, the *k*-mer Jaccard Similarity is intuitively still a good proxy for the overlap size in the presence of errors, as this expression is monotone increasing in the true alignment. The parameter *k* should be large enough to guarantee that not too many spurious *k*-mer collisions occur, but small enough so that a reasonable number of *k*-mers per read are not corrupted by noise [Myers, 2014, Berlin et al., 2015, Li, 2016]. For a relative noise rate of 30% (which results from both reads having error rates of around 15%), Myers [2014] argues that *k* = 7 achieves the optimal trade off. In the remainder of the manuscript, we utilize *k* = 7.

While Figure 3 depicts an *overlap* between *S*_0_ and *S*_1_ (i.e., a suffix of *S*_0_ that matches a prefix of *S*_1_), the general alignment problem is concerned with finding long matches between *S*_0_ and *S*_1_ which need not be proper overlaps. In general, one may think of *α*_0,1_ ∈ [0, 1] as the total size of the matches between *S*_0_ and *S*_1_, which may include an overlap and repeats. For simplicity, we will focus our discussion on overlaps, and show that this is a sufficiently good model to predict alignment well.

Computing the Jaccard Similarity between two reads of length *L* takes *O*(*L*) time. Hence computing this metric for all pairs of reads would take *O*(*n*^2^*L*) time, which is quite expensive. An attractive feature of the JS metric is that it can be efficiently estimated. A probabilistic approach for estimating JS through the use of *min-hashes* was originally proposed in Broder et al. [1997]. In essence, one takes a random hash function *h*, hashes all *k*-mers in a read *S*_*i*_ and picks the minimum hash value. Define

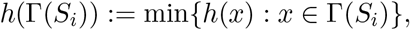

for some hash function *h* and read *S*_*i*_. Then we observe that, for a randomly chosen hash function *h*,

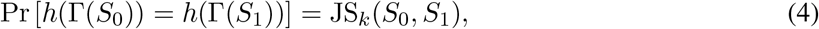

since *h* is equally likely to have any of the *k*-mers on both reads as its minimizer. This surprising fact is elaborated upon and proved in Section 3.3.3 of Leskovec et al. [2014]. This means that we can use a random hash function to get an unbiased estimate of the Jaccard similarity between two strings. More precisely, if we sample random hash functions *h*_1_, *h*_2_, …, *h*_*H*_, we can estimate the Jaccard Similarity as

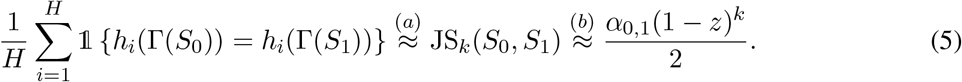

Hence, by choosing *H* moderately large, one should be able to accurately estimate JS_*k*_(*S*_0_, *S*_1_), which provides a proxy for the alignment size. With *H* hashes, one would take *O*(*nLH*) time to compute the hashes and *O*(*n*^2^*H*) time to compute collisions, which is *O*(*n*^2^*H*) time in regimes of interest where *n* ⨠ *L*.

### 2.1 Drawbacks of Jaccard Similarity

The key assumption that drives the approximation in equation (5b) is that that all *k*-mers are roughly equally likely to occur in the reads. On real datasets however, *k*-mer distributions are far from uniform, as illustrated in Fig. 4(b).

**Figure 4:**
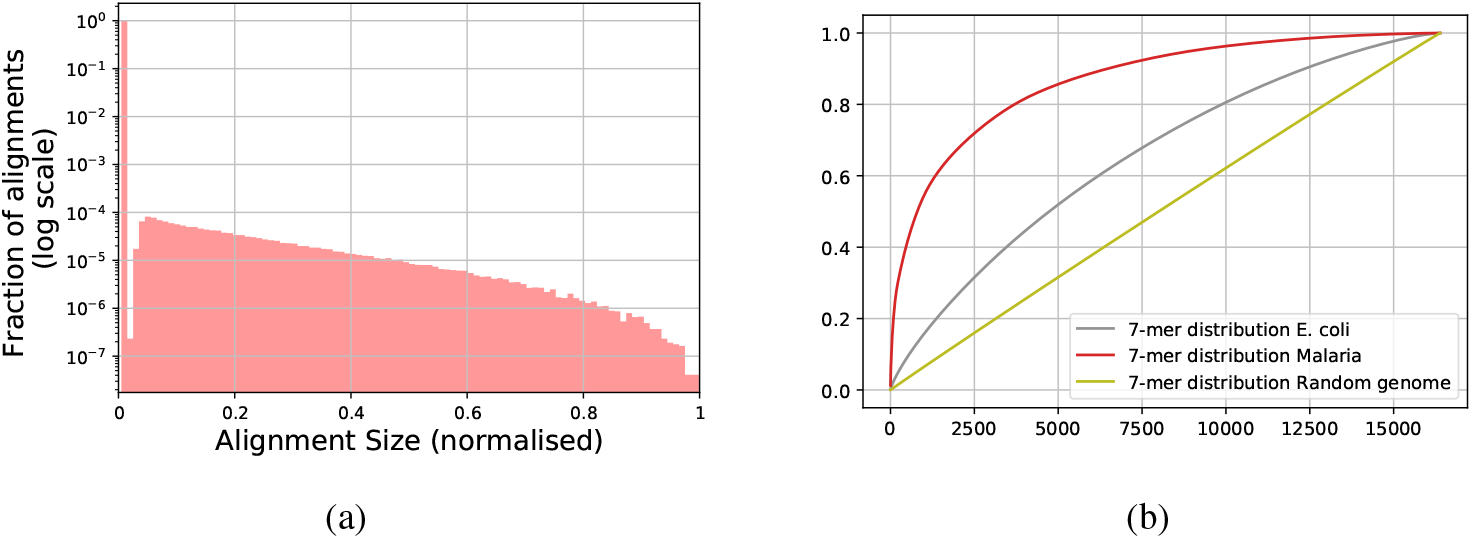
(a) A histogram of fraction of shared sequence detected by Daligner Myers [2014] in reads of *E. coli* K-12 dataset of Pacific Biosciences Inc. [2013]. We note that more than 99.9% of pairs of reads have no alignment between them. We also note that practical aligners are not able to capture small overlaps, which are hard to distinguish from spurious alignments generated by noise, creating the “notch” in the histogram. (b) Cumulative distribution function (CDF) of the *k*-mer distributions for various genomes. For each genome, we sort the *k*-mers in decreasing order of frequency to help with visualization. We see that the distributions deviate significantly from a uniform distribution (dark yellow line).

An uneven distribution of the *k*-mers throughout a genome increases the likelihood that the non-overlapping parts of two reads *S*_0_ and *S*_1_ share *k*-mers. In this case, for a randomly drawn hash function *h*, (4) still holds, but equation (5b) no longer holds. In particular, if the hash function *h* is such that common *k*-mers are given low hash values, the min-hash *collision probability*, given by Pr [*h*(Γ(*S*_0_)) = *h*(Γ(*S*_1_))], can be significantly higher than the right-most expression in equation (5). For this reason, when the *k*-mer distribution throughout a genome is uneven, (5) yields a poor estimate for the read overlap size. This is illustrated in Fig. S13(a) for *E. coli* reads from [Pacific Biosciences Inc., 2013], where we show that Jaccard similarity is a poor predictor of alignment sizes.

One simple way to address this issue is to “mask” common *k*-mers [Myers, 2014, Koren et al., 2017], and then compute the Jaccard similarity on the remaining *k*-mers. However, these approaches are arbitrary and require the tuning of parameters that can in general depend on the distribution. Intuitively, they can be thought of as applying a hard threshold to determine which *k*-mer matches are due to noise, and which are actual signal. Our approach can be thought of as soft version of this thresholding, where we weight the importance of each min-hash collision differently.

## 3 Spectral Jaccard Similarity

We propose a new Jaccard-similarity-inspired approach to estimate the overlap between reads that avoids the need for hard thresholds for determining “common *k*-mers” or “bad hashes” and instead assigns soft penalties to individual hash functions according to how biased an estimator they are for alignment size.

Suppose reads *S*_0_ and *S*_1_ of length *L* have an overlap of size *αL* for some 0 ≤ *α* ≤ 1, and no other significant repeats across them. If there were no shared *k*-mers in the non-overlapping part of the reads, we would model the min-hash collision event for a random hash function *h*, as

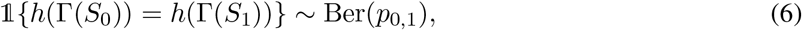

where 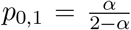 (this expression can be modified to account for errors as in (3)). However, when the distribution of *k*-mers is uneven, the min-hash collision probability is larger than *p*_0,1_. Moreover, some hash functions are worse than others: if *h* assigns lower values to common *k*-mers, it is more likely to overestimate *p*_0,1_. We model this effect by rewriting (6) as

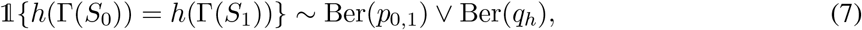

where ⋁ is the boolean “or” operator and *q*_*h*_ ∈ [0, 1] is a hash-specific parameter that can be intuitively understood as how unreliable *h* is due to common *k*-mers. Notice that the hash-specific noise term always leads to overestimation of *p*_0,1_. We also emphasize that the *q*_*h*_’s are unknown. Therefore, we cannot directly estimate the *p*_*i,j*_ and instead we need to jointly estimate all model parameters.

In order to perform this joint estimation, we define the *min-hash collision matrix* as follows. For a fixed reference read *S*_0_, a list of target reads *S*_1_, *S*_2_, …, *S*_*n*_, and a list of hash functions *h*_1_, …, *h*_*H*_, the (*i, j*)th entry of the min-hash collision matrix is the binary indicator variable for whether there is a min-hash collision between *S*_0_ and *S*_*i*_ when using hash function *h*_*j*_; i.e., 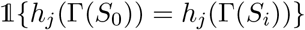. Notice that JS_*k*_(*S*_0_, *S*_*i*_) can be directly estimated from the min-hash collision matrix by computing the fraction of 1s in the *i*th row.

As it turns out, if we assume that the entries in the min-hash collision matrix were generated according to (7), we can jointly estimate the *p*_0,*i*_’s and the 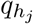’s by performing a singular value decomposition (SVD) on an offset version of the min-hash collision matrix. This allows us to use efficient algorithms for computing the SVD in order to obtain estimates for the parameters *p*_0,*i*_.

We refer to the parameters *p*_0,*i*_ as the Spectral Jaccard Similarity (SJS) between *S*_0_ and *S*_*i*_. Intuitively, this value can be understood as a “discounted” version of JS_*k*_(*S*_0_, *S*_*i*_), where we discount the contribution of common *k*-mers. It is important to point out, however, that *p*_0,*i*_’s are only implicitly defined as model parameters that are learned from the data, and no explicit formula for them exists. Next we describe the computation of the SJS in more detail.

### 3.1 Computing the Spectral Jaccard similarity

Algorithmically, we approximate all pairwise alignments by iterating over each read in the dataset, treating it as the reference read, and computing the SJS between the reference and all other reads. For a reference read *S*_0_, we define the min-hash collision matrix as *A*_0_ ∈ {0, 1}^*n*×*H*^ where

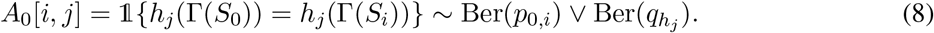

For cleanliness of notation, we will write *p*_0,*i*_ = *p*_*i*_ and 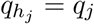. Notice that both the *p*_*i*_’s and the *q*_*j*_’s depend on the choice of *S*_0_, but we do not make that dependence explicit in the notation.

The key observation about our model is that, in expectation, the matrix *A*_0_ defined in (8) is rank one after accounting for offset. More precisely, since 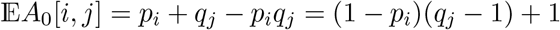 we have

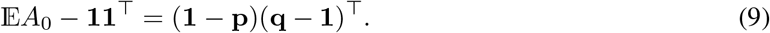

We illustrate this point by comparing the sorted singular values of *A*_*i*_ − 11^⊤^ for the PacBio *E. coli* K-12 dataset, shown in Figure S14. Focusing on read 0, we note that the fact that *A*_0_ − 11^⊤^ is in expectation a rank-one matrix, allows us to estimate p and q through an SVD on *A*_0_ − 11^⊤^. More precisely, if we let u and v be respectively the leading left and right singular vectors of *A*_0_ − 11^⊤^, then we expect u to be approximately proportional to (1 − p) and v to be approximately proportional to (q − 1), up to flipping signs. We then normalize *q*_*j*_’s to be between 0 and 1, assuming that min_*j*_ *q*_*j*_ = 0. For *p*_*i*_’s, we require a slightly more sophisticated normalization method, which we discuss in Appendix A.

To illustrate the comparison between SJ and SJS we consider the example shown in Figure 5. The standard min-hash JS approach would estimate JS_*k*_(*S*_0_, *S*_*i*_) to be the fraction of 1s in the *i*th row. We see that while rows 1 and 3 have the same estimated JS, they have different SJS values. This is because columns 2 and 5 are found to be noisier (i.e., worse hash functions), and so while rows 1 and 3 have two collisions each, a collision on column 1 is deemed more indicative of alignment, and thus row 3 has a higher SJS than row 1. As it turns out, the estimates of *p*_*i*_ obtained via SVD are a much better proxy for the size of the alignment between reads *S*_0_ and *S*_*i*_ than the standard Jaccard similarity, particularly when the *k*-mer distribution is uneven. To illustrate this point, we computed the SJS values *p*_*i*_ (from a min-hash collision matrix with *n* = 1004 and *H* = 1000) and the *exact* Jaccard Similarity JS_*k*_(*S*_0_, *S*_*i*_) for the corresponding pairs of reads, for the *E. coli* PacBio dataset. As illustrated in Fig. S13, while the *R*^2^ coefficient between (exact) JS and overlap size is only 0.18, for SJS the *R*^2^ coefficient is 0.48.

**Figure 5:**
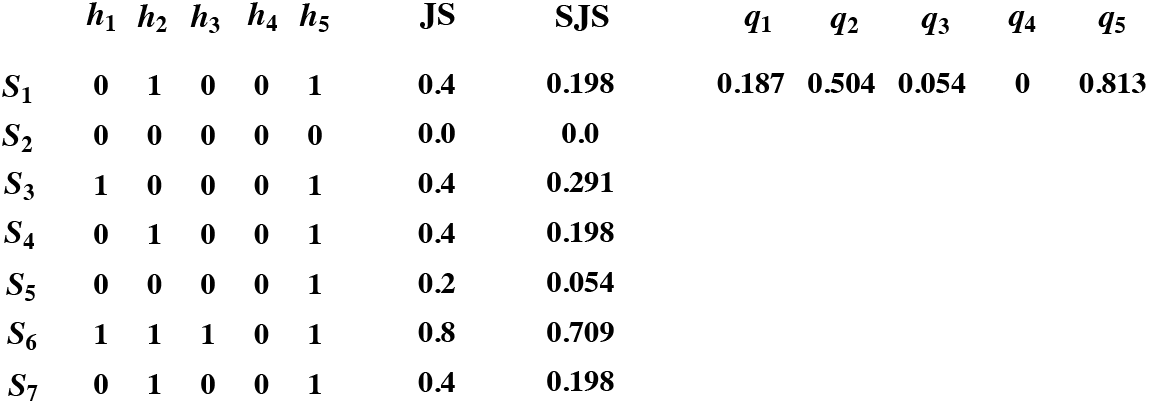
Example of comparison between JS and SJS on a small matrix. While the standard JS approach would assign the same value to rows 1 and 3, SJS takes into account the fact that columns 2 and 5 are seen as less reliable indicators of alignment.

While *p*_*i*_ parameters, or the SJS values, track the alignment between pairs of reads, we have not given a precise meaning to the *q*_*j*_s we recover. Intuitively, a large *q*_*j*_ means that the corresponding hash function is more likely to create a spurious min-hash collision, thus being a less reliable estimator for the alignment size. We provide a more in-depth interpretation in Section 5.2. In Figures 9(a) and 9(b), we plot the frequency of the argmin *k*-mer for different hash functions, and verify that large *q*_*j*_s correspond to hash functions whose argmin *k*-mers have high frequency. We point out that we compute SJS on a reference-by-reference basis instead of considering a single matrix with all 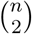 rows at once. While SJS can be computed on the matrix with all 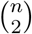 read pairs as rows, allowing the *q*_*j*_ hash parameters to be reference-specific increases their ability to capture the discriminative power of each hash function, as this depends on which *k*-mers are present in the reference read. Furthermore, it avoids the computation of an SVD for a very large matrix.

From a computational perspective, this algorithm computes all *H* hashes in *O*(*nLH*) time, collisions in *O*(*n*^2^*H*) time, and performs *n* (*n* × *H*) SVDs. In regimes of interest where *n* ⨠ *L* this last term dominates. Computing a full (*n* × *H*) SVD requires *O*(min(*n*^2^*H, nH*^2^)) = *O*(*nH*^2^), giving a computational complexity of *O*(*n*^2^*H*^2^), which naively is slower than the min-hash based JS computation. However, note that we only need to compute the principal left singular vector, which can be done efficiently via power-iteration, reducing he *O*(*nH*^2^) run time to 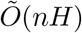, where 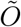 suppresses logarithmic factors in *n* and *H*. Further improvement is available in the practical case whenever most of the (*S*_0_, *S*_*i*_) pairs have no overlap, as most of the *p*_*i*_’s are expected to be close to zero. When this holds true, we are able to approximate the principal right singular vector q efficiently, allowing us to approximate p via a single matrix-vector product, speeding up our method significantly to *O*(*n*^2^*H*), the same complexity of JS. This approach is discussed in more detail in Section 5.1.

## 4 Results

To compare the performance of JS and SJS at estimating alignment sizes we focus on two PacBio datasets (an *E. coli* dataset [Pacific Biosciences Inc., 2013] and a *K. pneumoniae* (NCTC5047) dataset [Parkhill et al]) and consider the problem of identifying pairs of reads with an overlap of size at least *θ*. We compute exact JS values and compare them with SJS values computed based on 1000 hash functions (see Section 5.3 for results with different numbers of hash functions). We discuss the preprocessing steps performed on these datasets in Appendix C. We utilize Daligner [Myers, 2014] alignments as ground truth for the alignment sizes. In Figure 6, we plot ROC curves for different values of *θ*. We point out that using the Daligner outputs as ground truth is not ideal, since the tool itself utilizes an empirical Jaccard Similarity based filter to align reads; this choice of ground truth biases the result in favor of the conventional Jaccard similarity. Despite this we note that SJS performs significantly better than JS on all data sets tested, even when the output metric inherently favors JS. In addition, we obtain similar results when Minimap2 is used instead of Daligner (Figure S12).

**Figure 6:**
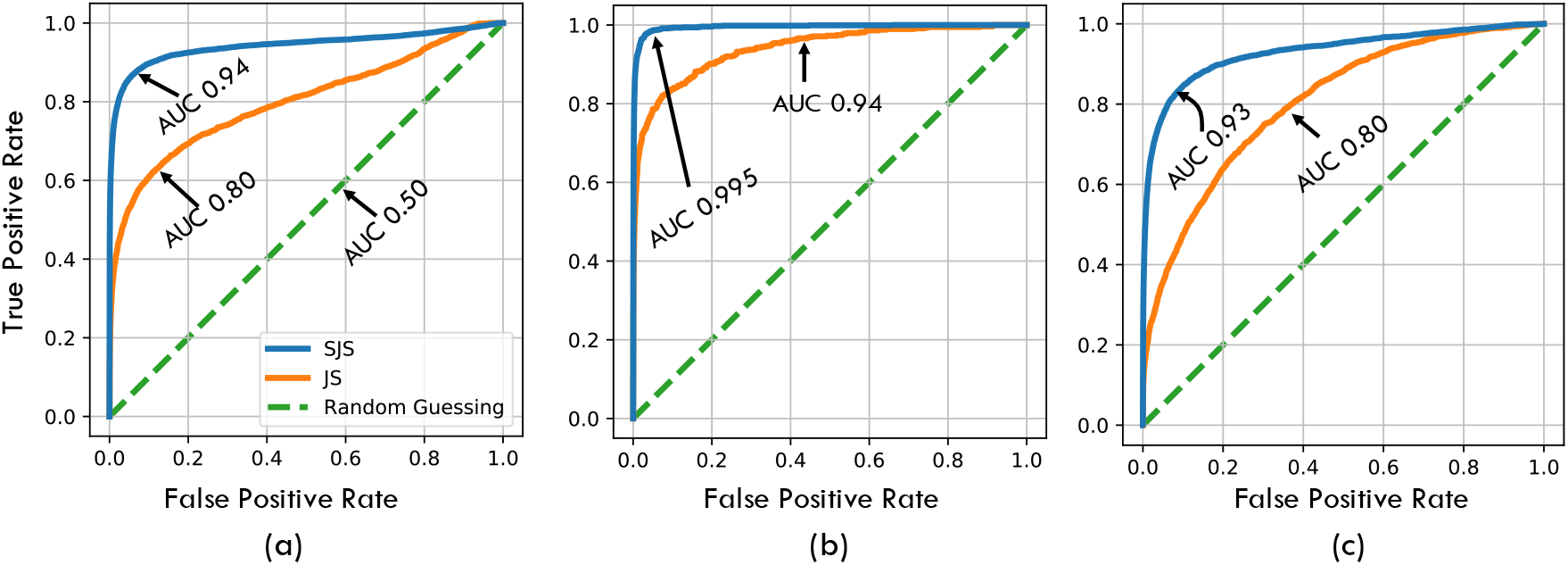
ROC curves across different PacBio datasets and different *θ* thresholds using Daligner ground truth and 1000 hashes: (*a*) shows ROC of *E. coli* (K-12 from PacBio website) for alignment threshold *θ* = 0.3, (*b*) shows ROC of *E. coli* for alignment threshold *θ* = 0.8, and (*c*) shows ROC of *K. Pneumoniae* (NCTC5047) for alignment threshold *θ* = 0.8. Figure S15 shows a similar plot with minimap2 ground truth.

Minimap2 and Daligner use different procedures to filter pairs of reads. This provides additional evidence that the superior performance of SJS over JS is not due to using Daligner to define the ground truth.

We note that the performance of both JS and SJS filters degrades as the *k*-mer distribution becomes skewed. To formally capture this skew for a dataset of reads *D* = *S*_1_, …, *S*_*n*_, each of which is assumed to have length *L*, we let the *k*-mer distribution of *D* be defined as the empirical distribution of the *n*(*L* − *k* + 1) *k*-mers in the reads of *D*. To formally capture this skew for a dataset of reads *D* = *S*_1_, …, *S*_*n*_ with an average read length of *L*, we let the *k*-mer distribution of *D* be defined as the empirical distribution of the ≈ *nL k*-mers in the reads of *D*. We then have the following definitions.

### Definition 1.

*For a dataset D and hash function h, the collision probability of read S, denoted* colprob_*D,h*_(*S*), *is the probability that a set of L* − *k* + 1 *randomly chosen k-mers drawn independently and uniformly at random from the k-mer distribution of D has the same min-hash on hash h as S.*

This collision probability of read *S* in a dataset *D* with hash *h* can be computed in closed form, as discussed in Appendix B. Further, we can extend this definition in order to capture the overall hardness of approximating pairwise alignments in a dataset *D* as follows.

### Definition 2.

*The mean collision probability of a dataset D* = {*S*_1_, …, *S*_*n*_} *is given by*

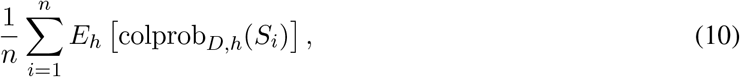

*where E*_*h*_ *is the expectation with respect to a randomly chosen hash function h*.

While computing the expectation *E*_*h*_ over hash functions *h* is computationally infeasible, it can be approximated by an average over a set of randomly chosen hash functions *h*_1_, …, *h*_*m*_. As we discuss in Appendix B, we can use the set of hash functions that were used to compute the min-hashes to give a closed form approximation of the mean collision probability of a dataset.

In Fig. 7(a) we plot the performance of SJS and JS as a function of the computed collision probabilities for 40 datasets from the NCTC3000 project [Parkhill et al]. This shows the uniform improvement in performance afforded by SJS, in that for every dataset the SJS AUC is higher than the JS AUC. Further, it shows that as the *k*-mer distribution becomes more skewed, the degradation in performance suffered by SJS is smaller than that suffered by JS. We plot the ratio of the improvement of the two AUCs over random guessing in Fig. 7(b). This shows that the improvement of SJS over JS is larger when the *k*-mer distribution is more skewed.

**Figure 7:**
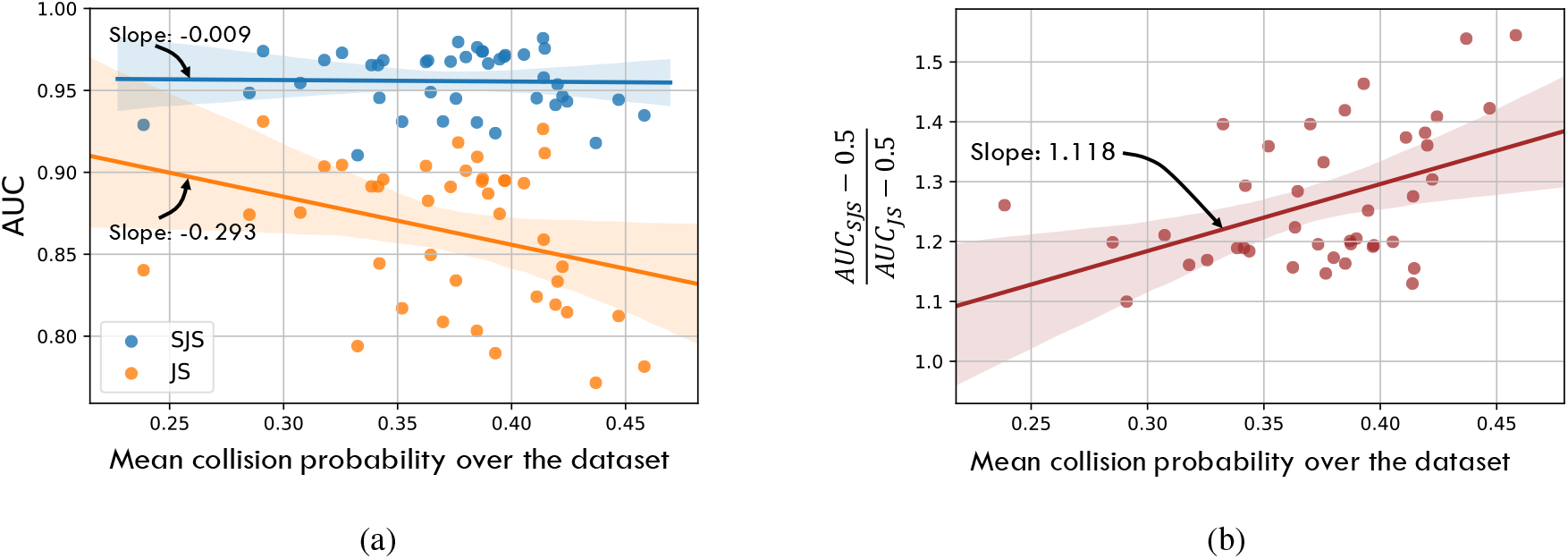
(a) The higher the min-hash collision probability is, the worse both methods perform, indicating a “harder” dataset. However, the performance of the SJS filter degrades less than that of the JS filter; (b) Ratio between the improvement of the SJS filter over random guessing and the improvement of the JS filter over random guessing, as a function of collision probability of the reads’ *k*-mer distribution. The results in both plots were computed for *θ* = .3 using 1000 hashes.

## 5 Discussion

In this paper, we introduced the notion of Spectral Jaccard Similarity as an alternative to the standard *k*-mer Jaccard Similarity for estimating the overlap size between pairs of noisy, third-generation sequencing reads. SJS is a probabilistic approach that utilizes min-hash collisions as a way to estimate the size of the overlap between pairs of reads. However, unlike previous approaches, SJS attempts to learn how good different hash functions are at estimating overlap size for that *specific dataset*. In particular, when the *k*-mer distribution of the dataset in question is very uneven, the gain of SJS over JS is greater.

We conclude the paper by discussing some additional aspects of the algorithm implementation and providing some further validation of the model. First, we show how the fact that the columns of the *A*_0_ matrix are typically sparse can be exploited in order to approximate the SVD using a single matrix-vector multiplication, which can significantly speed up the computation of SJS. Second, we validate our earlier claim that *q*_*j*_s represent how bad a hash function is for the purpose of alignment. Third, we examine the performance of SJS as a function of the number of hashes used and show that it can match the performance of exact Jaccard similarity with around 150 hashes.

A final implementation-related point, the calibration of the SJS values across different reads, is discussed in Appendix A. More precisely, we describe how we normalize the *p*_*i*_s obtained for different reference reads (i.e., from the SVD of different matrices *A*_*i*_ and *A*_*j*_) so that the SJS values are comparable.

### 5.1 Approximation in the case where most *p*_*i*_’s are zero

Given a min-hash collision matrix *A* ∈ {0, 1}^*n*×*H*^, define

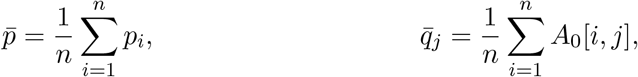

for 1 ≤ *j* ≤ *H*; i.e., 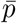 is the average *p*_*i*_ value and 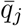 is the fraction of ones in column j. We note that, since *A*_0_[*i, j*] ~ Ber(*p*_*i*_) ⋁ Ber(*q*_*j*_), when most *p*_*i*_’s are zero, most of the entries in column *j* are distributed as Ber(*q*_*j*_). It follows that 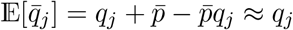 since 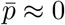. This means that the leading right singular vector is approximately 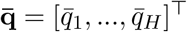. Since the rank-one approximation is

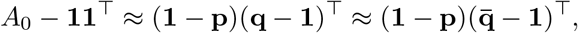

by multiplying both sides by 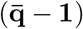, we obtain

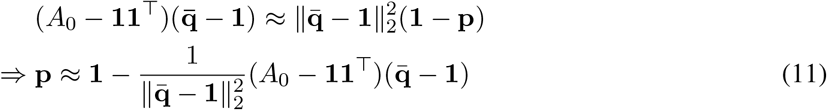

which gives us a method to compute the Spectral Jaccard Similarity with a matrix-vector multiplication, rather than an SVD. This can intuitively be understood as follows. We wish to approximate the principal left singular vector of the matrix *A*_0_ − 11^⊤^. We are, however, given some side information; we are able to easily obtain a high-quality approximation of the principal right singular vector as 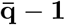. This allows us to effectively perform one step of the Power Iteration algorithm, as 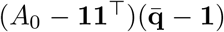, which after normalization, gives us a very good approximation of the principal left singular vector.

In Figure 8(a), we show that for an *E. coli* dataset where most reads do not have any overlap, 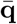 is very correlated with q. In Figure 8(b), we show that the approximate SJS values computed using (11) are highly correlated with those computed through a full SVD.

**Figure 8:**
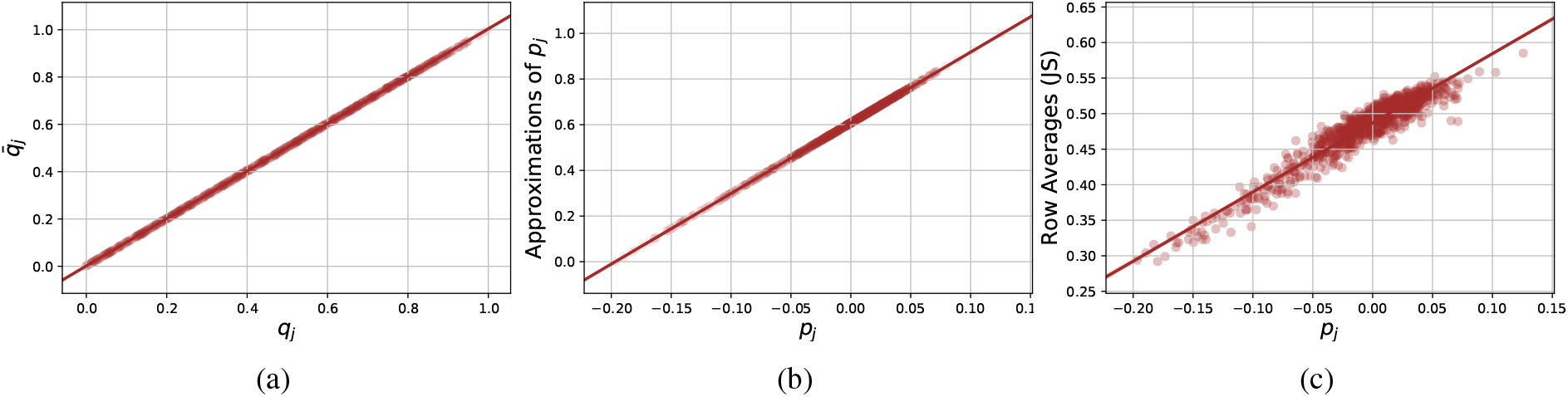
When most *p*_*i*_ ≈ 0, it is possible to approximate the SVD by a simple matrix-vector multiplication as described in (11). In particular, we verify empirically that (a) 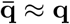 and that (b) the p obtained from (11) is nearly the same as the one computed by SVD. (c) If one instead tries to approximate p by considering row averages, the approximation is not as good.

**Figure 9:**
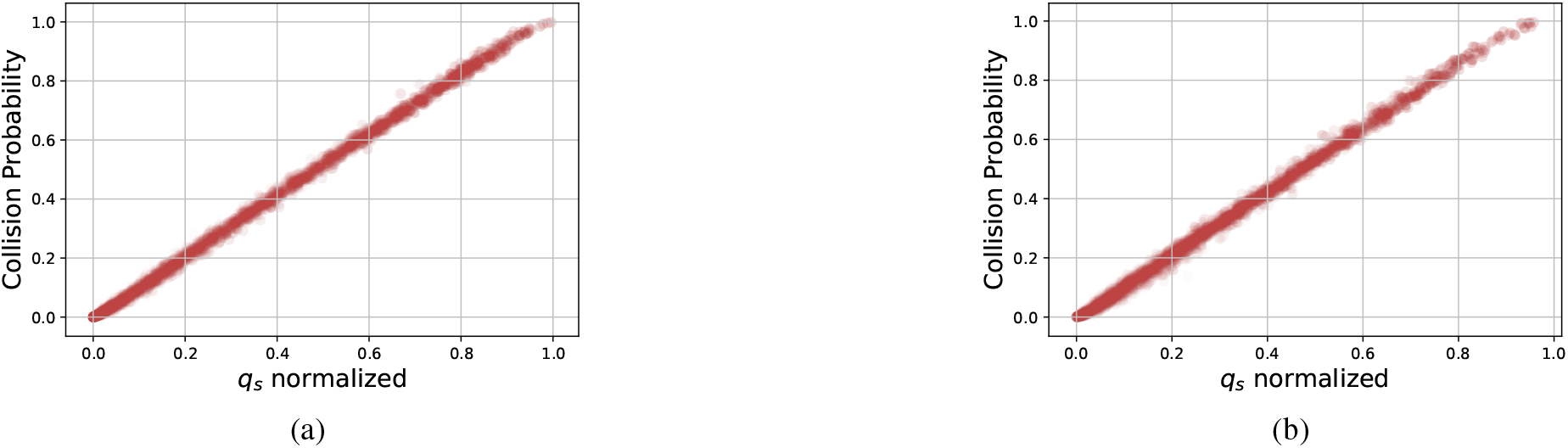
For each hash function *h*_*j*_, we compared the collision probability on hash *h*_*j*_ with the corresponding *q*_*j*_ for (a) the *E. coli* dataset and (b) the *K. pneumoniae* dataset (NCTC5047).

We note that one could have have attempted to use row averages instead of column averages in this approximation procedure. However, this would correspond to computing the standard Jaccard Similarities. Jaccard Similarity is not as well correlated with the Spectral Jaccard Similarity as we show in Figure 8(c). Further, we note that (11) implies that 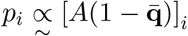, which is expanded as

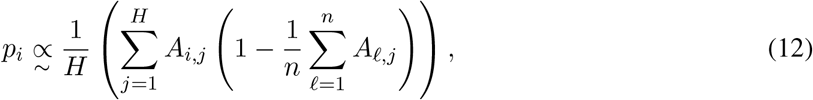

where 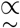 indicates “monotonic function of”. Since 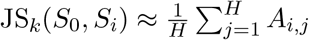, our method can be understood as down-weighting the contribution of hash functions that yield many collisions.

We show that this approximation performance performs nearly as well as SJS in Figure S10.

### 5.2 Validating the Model

In Section 3, we proposed the model in (7) with the interpretation that *q*_*j*_ represented how likely were min-hash collisions given the hash function *h*_*j*_. In this section, we empirically verify that claim. In Figure 9, we show the collision probability of a reference read on a hash function *h*_*j*_ as a function of our computed *q*_*j*_ for the *E. coli* and *K. pneumoneae* (NCTC5047) datasets. We see a very strong correlation between the computed collision probability and the *q*_*j*_ parameters, validating our model.

### 5.3 Trade-off between filter accuracy and number of hash functions

While throughout this manuscript we present results for SJS using 1000 hash functions, our method performs well even with a smaller number of hash functions. In Figure S17(a), we plot ROC curves for SJS and the SJS approximation (aSJS) described in Section 5.1 using different number of hashes in order to compare the performance of these filters on the *E. coli* K-12 dataset. We note that as few as 150 hashes are enough for SJS to dominate the exact JS based filter. The performance of the approximation is similar.

**Table 1:**
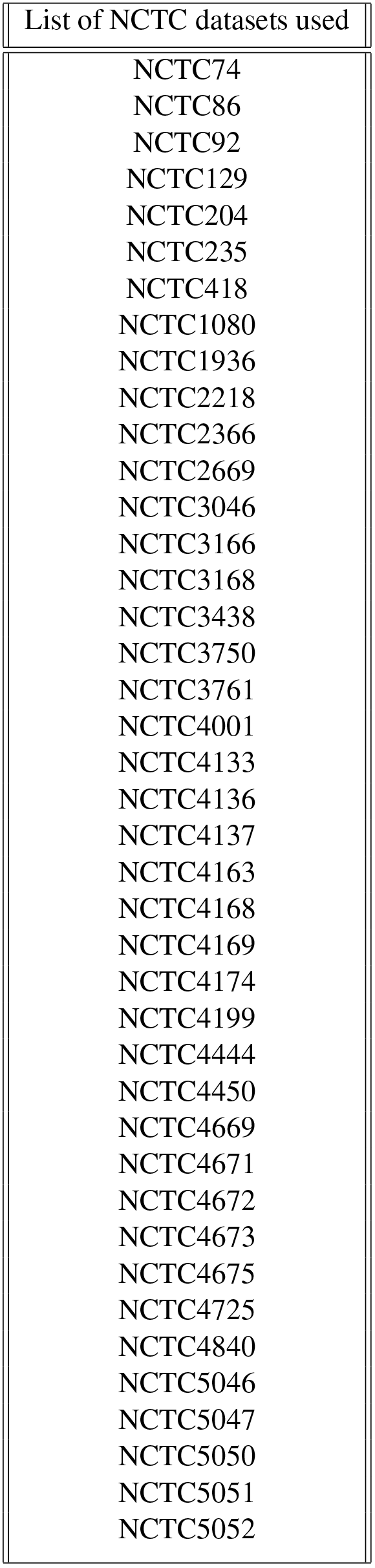
List of datasets used

## 6 Acknowledgements

The authors gratefully acknowledge funding from the NSF GRFP, Alcatel-Lucent Stanford Graduate Fellowship, NSF grant under CCF-1563098, and the Center for Science of Information (CSoI), an NSF Science and Technology Center under grant agreement CCF-0939370.

## Appendices

## A Calibrating SJS values from different reference reads

Given two min hash collision matrices *A*_0_, *A*_1_ ∈ {0, 1}^*n*×*H*^ we have from (9) that *A*_*k*_ ∈ 11^⊤^ ≈ (1 − p_*k*_)(q_*k*_ − 1)^⊤^ for *k* ∈ {0, 1}. When one computes an SVD, the left and right singular vectors are scaled so that they have an *ℓ*_2_ norm of 1. However for our purposes, normalizing these vectors by the *ℓ*_2_ norm implies assuming that p_*i*_’s have similar norms for all reference reads *i* which need not be true. Notice that we could potentially have two constants *γ*_0_, *γ*_1_ ≠ 0, *γ*_0_ ≠ *γ*_1_ such that 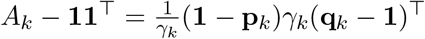 for *k* ∈ {0, 1} With different *γ*_0_ and *γ*_1_, we see that we would not be able to compare values in p _0_ and p _1_, which leads to issues when one is trying to threshold all with a common threshold.

One approach to calibrating the *p*_0,*i*_’s and *p*_1,*i*_’s to make the left singular vectors of *A*_0_ − 11^⊤^ and *A*_1_ − 11^⊤^ comparable takes advantage of *calibration reads*. Calibration reads are fake reads, following the dataset’s *k*-mer distribution, that are expected to have no overlap with the reference, thus making sure that the Spectral Jaccard Similarities computed on different runs of the SVD have the same “ground level”, so that we can compare the SJS values between two pairs of reads coming from two different SVDs.

More precisely, our procedure involves including *W* calibration reads into the min-hash collision matrix. To create these calibration reads, we sample *L* − *k* + 1 *k*-mers from the *k*-mer distribution of the dataset, where *L* is the average read length of the dataset. Hence, each calibration read is essentially a “bag of *L* − *k* + 1 *k*-mers”. Then we augment the matrix *A*_0_ and consider instead a matrix *B*_0_ ∈ {0, 1}^(*n*+*W*)×*H*^ with the first *n* rows of *B*_0_ being *A*_0_ and the last *W* rows being the min-hash collision matrix computed for the *W* calibration reads. We then compute an SVD of *B*_0_ − 11^⊤^ as

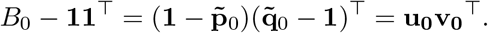

Using our random reads we define 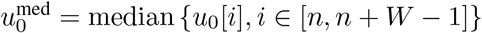. We now proceed to enforce that the median of 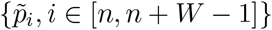 be 0 by defining the normalized SJS values as

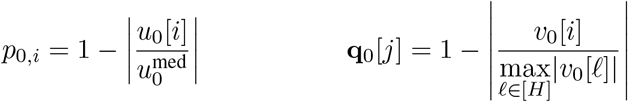

### Remark 1.

*We set the median alignment of the random reads to* 0 *rather than the minimum or maximum alignment because the variance in the extremal value was empirically found to be large.*

### Remark 2.

*In practice we use W* = 5, *but varying W does not seem to affect results significantly. Since n is typically large (in the thousands), these additional fake reads do not really change the result of the SVD.*

## B Collision Probability

In this section we describe how the collision probability (see Definitions 1 and 2) can be computed. We begin with some notation. First, note that each hash function can be thought of as inducing an ordering on the set of 4^*k*^ *k*-mers. We denote this permutation by (·)_*j*_ for hash *h*_*j*_. Let 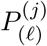 be the probability (or the frequency) of the *ℓ*-th *k*-mer in the permutation defined by hash *h*_*j*_ (on the *k*-mer distribution) of the dataset. For example, 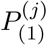 denotes the probability of the *k*-mer hashed to the minimum value by hash *h*_*j*_. Further, define partial sums

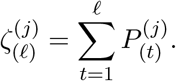

In words, 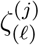 are the probabilities of the first *ℓ k*-mers in the permutation induced by hash *h*_*j*_ (or equivalently, the probability of *ℓ k*-mers that hash *h*_*j*_ maps to the smallest values). Finally, we denote the smallest *k*-mer in the order induced by hash *h*_*j*_ that is also in read *S*_*i*_ as *ℓ*_*i*_. Then the collision probability with the *i*th read as reference and hash function *h*_*j*_ is

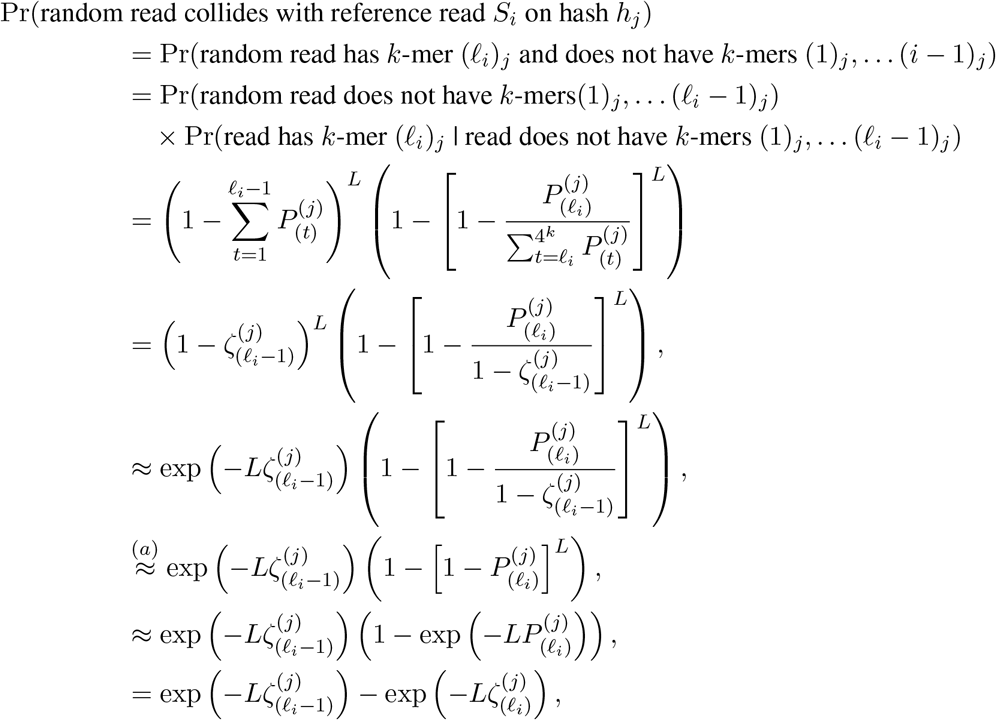

where the approximation in (*a*) follows in our setting where *L* and 4^*k*^ are of similar orders, and hence *ℓ*_*i*_ is small and 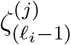 is close to 0. From this we obtain

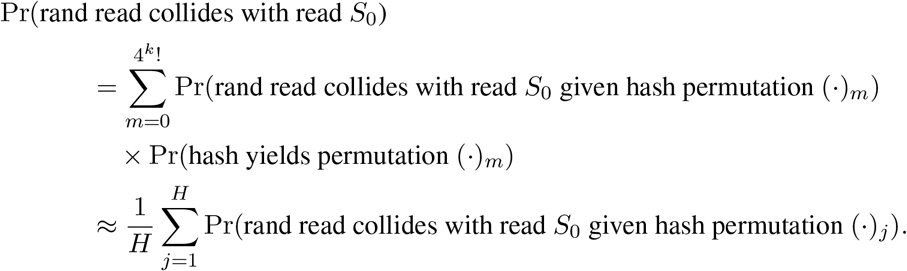

The first term above is exactly the probability we computed previously, while the second term is constant 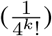. This gives us an explicit way to (approximately) compute the collision probability.

## C Details of experiments

We used the first 40 NCTC datasets of Parkhill et al (sorted numerically) for our empirical results. Since our main goal was to evaluate the performance of SJS (rather than fully re-align all the reads in these datasets), we pruned the datasets down so that each one contained 1000 reads. These reads were picked to match the distribution in Figure 4(a), such that each read has at least five significant overlaps. In essence, we went over the reads and added five reads that have large overlaps (according to Daligner [Myers, 2014]) with the read giving us a dataset where ~ 0.5% of read pairs have significant alignments. We then used Daligner and Minimap2 [Li, 2018] to generate alignments on these 1000 reads which we use as ground truths for AUC computations. The same is done for the *E. coli* K-12 dataset of Pacific Biosciences Inc. [2013] for the running example used throughout this manuscript. To obtain the ROC curves displayed we generated the binary indicator of whether each pair of reads had ground truth alignment above *θ*, then computed JS and SJS for each pair, and computed the ROC curve by considering different thresholds on the Jaccard and Spectral Jaccard similarities. The code to reproduce our numerical results is available online at https://github.com/TavorB/spectral_jaccard_similarity.

## D Additional Empirical Results

**Figure S10:**
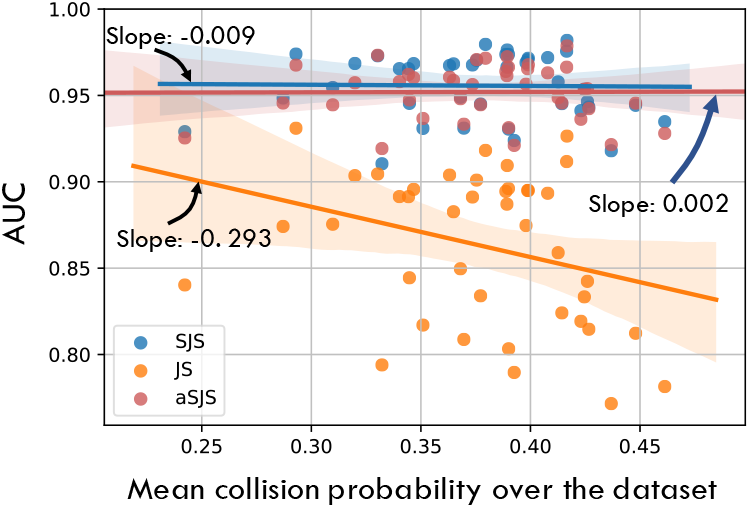
Redrawing Figure 7(a) with our power iteration approximation added in red as aSJS. As we can see, our one step of power iteration performs very well as an approximation for the SVD. While aSJS rarely outperforms SJS, it has most of the gain in predictive power while still being almost as computationally efficient as empirical JS, who’s performance is slightly under that of JS.

**Figure S11:**
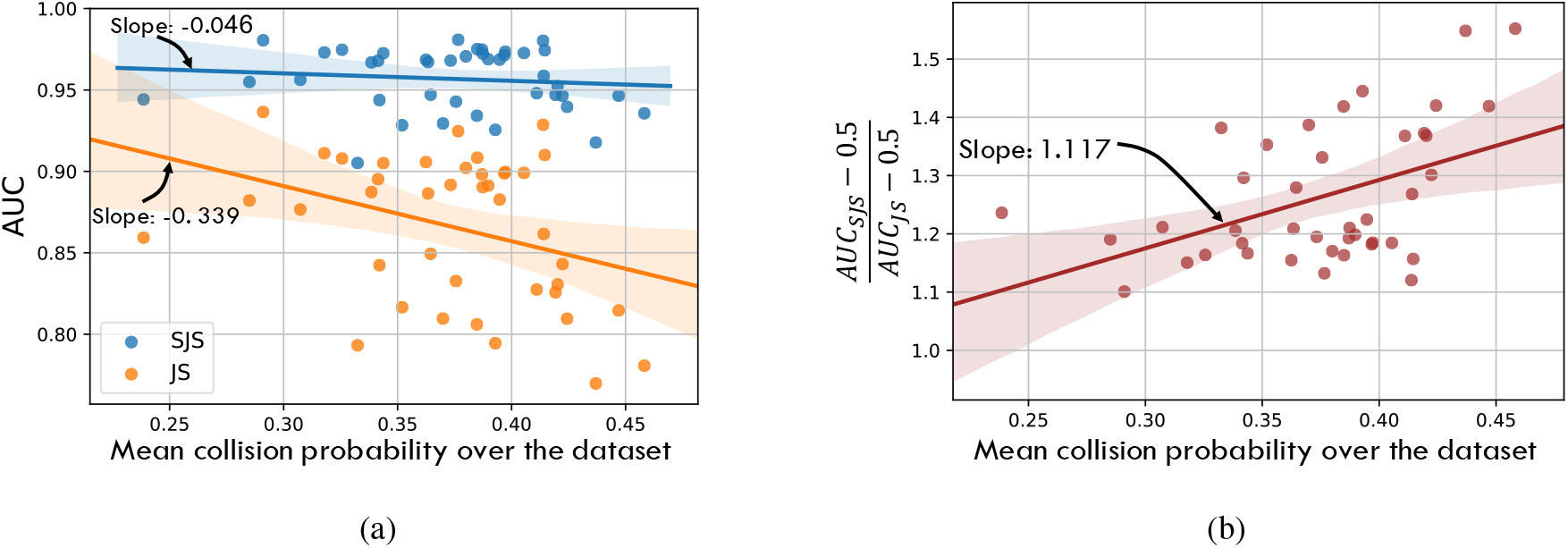
Analogue of Figure 7 using Minimap2 to provide the ground truth alignment sizes. The results are similar to those in Figure 7. Note that in (*b*) the slope is almost identical to what we obtained in the Daligner case in Fig. 7(b).

**Figure S12:**
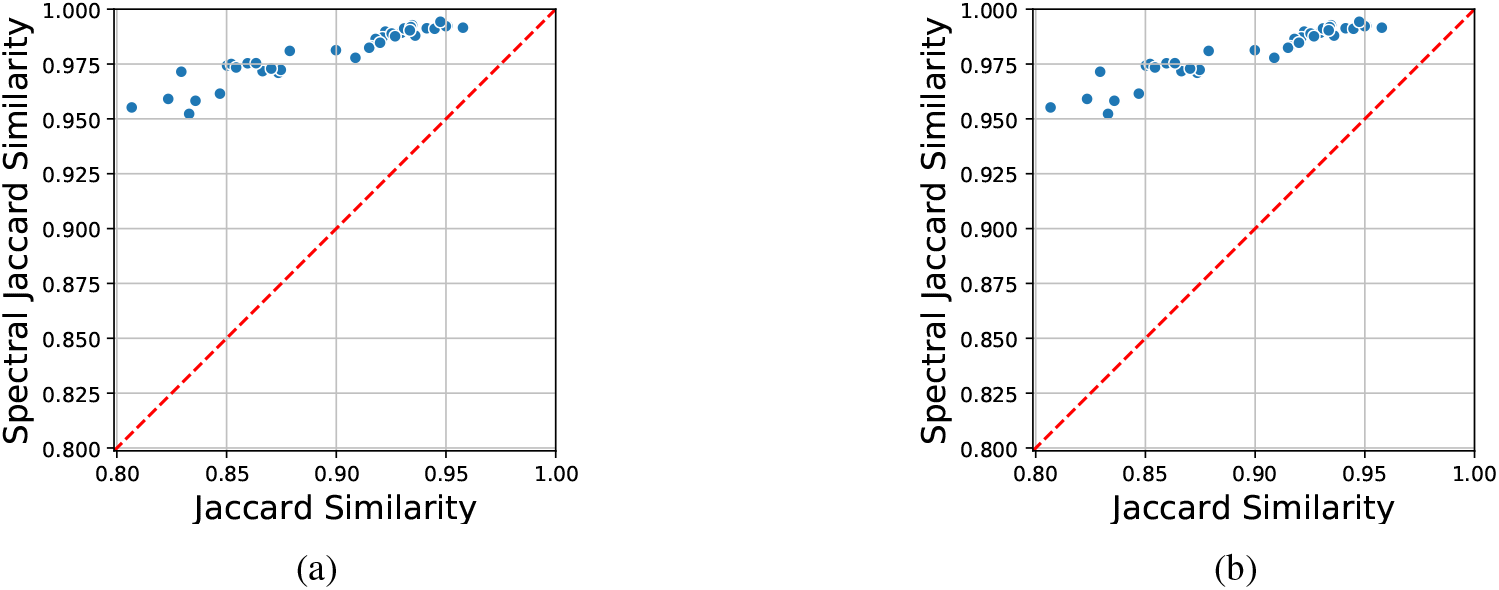
Analogue of Figure 2 but for *θ* = 0.5. (*a*) shows the ROCs using Daligner alignments as ground truth. (*b*) shows them using Minimap2 alignments as ground truth.

**Figure S13:**
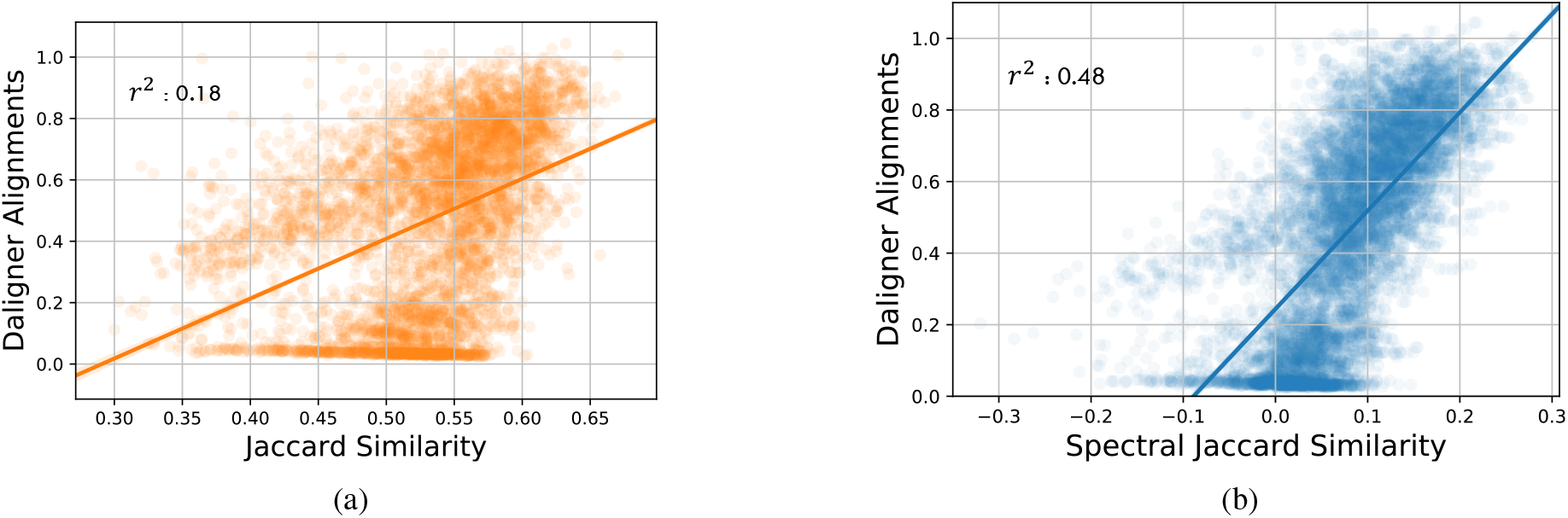
Linear regression fit to positive alignments found by Daligner to (*a*) Jaccard similarity between corresponding reads and (*b*) Spectral Jaccard similarity between the reads. Note that SJS provides a better fit.

**Figure S14:**
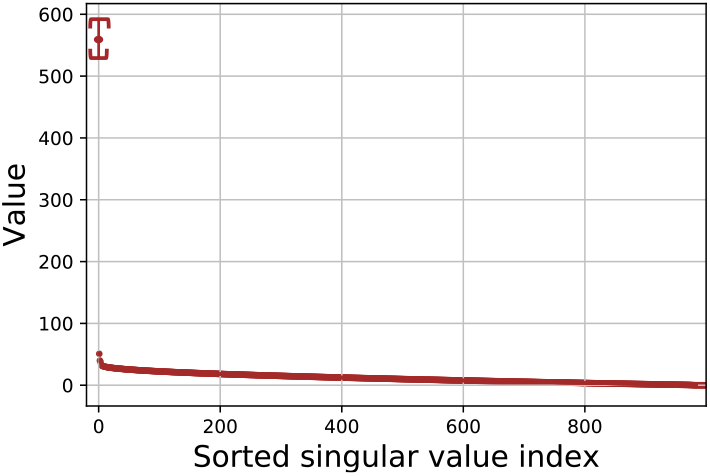
Singular values of empirical centered min-hash matrix *A*_*i*_ − 11^⊤^ for PacBio *E. coli* K-12 dataset. By repeating the computation of the singular values of *A*_*i*_ − 11^⊤^ for *i* = 1, 2, 3, *…* (i.e., based on different reference reads), we can compute the mean and standard deviation for each sorted singular value, which we display here using error bars of 1*σ*.

**Figure S15:**
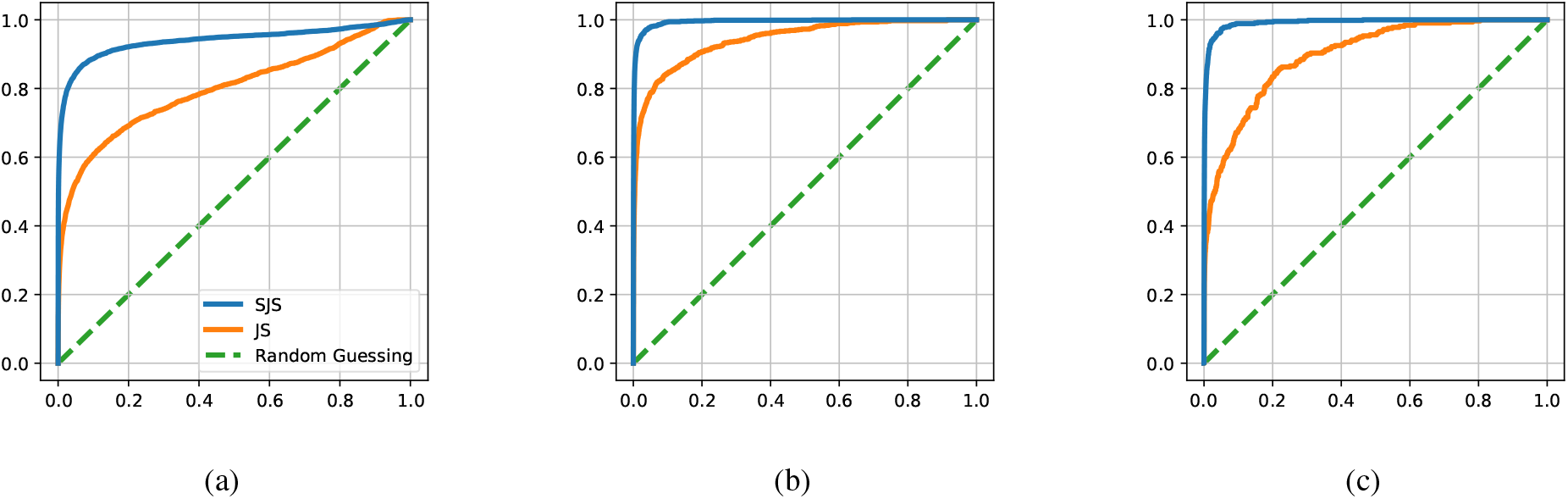
Analogue of Figure 6 but with minimap2 as ground truth alignments instead of Daligner. We plot ROC curves across different PacBio datasets and different *θ* thresholds: (*a*) shows ROC of *E. coli* (K-12 from PacBio website) for alignment threshold *θ* = 0.3, (*b*) shows ROC of *E. coli* for alignment threshold *θ* = 0.8, and (*c*) shows ROC of *K. Pneumoniae* (NCTC5047) for alignment threshold *θ* = 0.8

**Figure S16:**
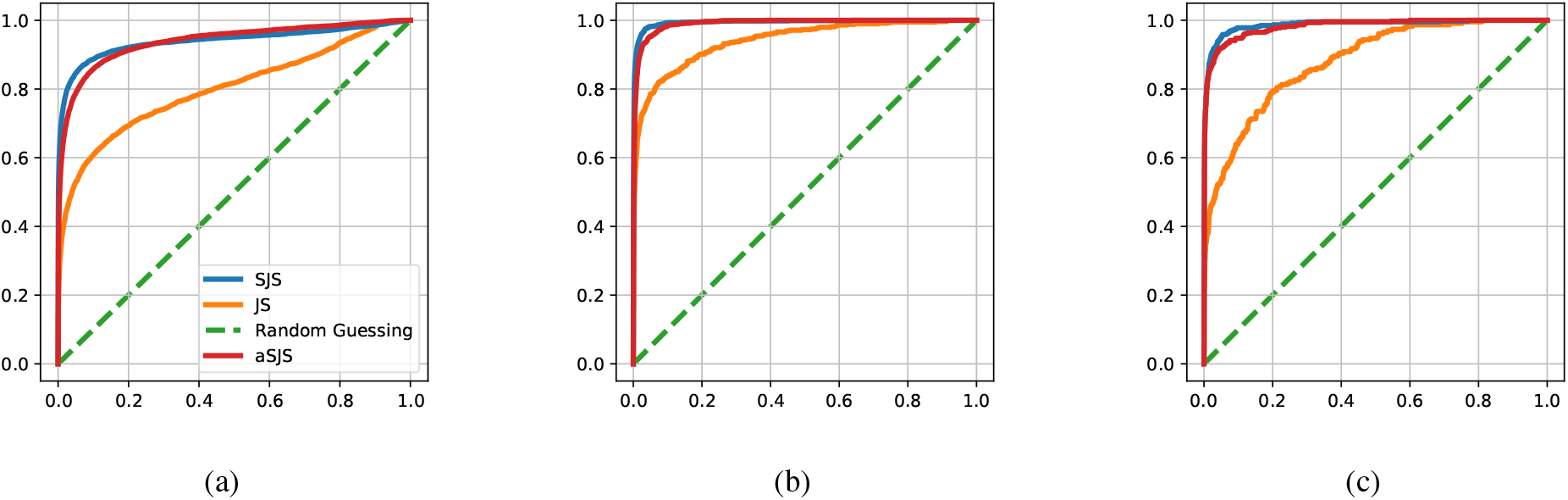
Analogue of Figure 6 with aSJS added (one step of power iteration approximation to SJS) on Daligner. We plot ROC curves across different PacBio datasets and different *θ* thresholds: (*a*) shows ROC of *E. coli* (K-12 from PacBio website) for alignment threshold *θ* = 0.3, (*b*) shows ROC of *E. coli* for alignment threshold *θ* = 0.8, and (*c*) shows ROC of *K. Pneumoniae* (NCTC5047) for alignment threshold *θ* = 0.8

**Figure S17:**
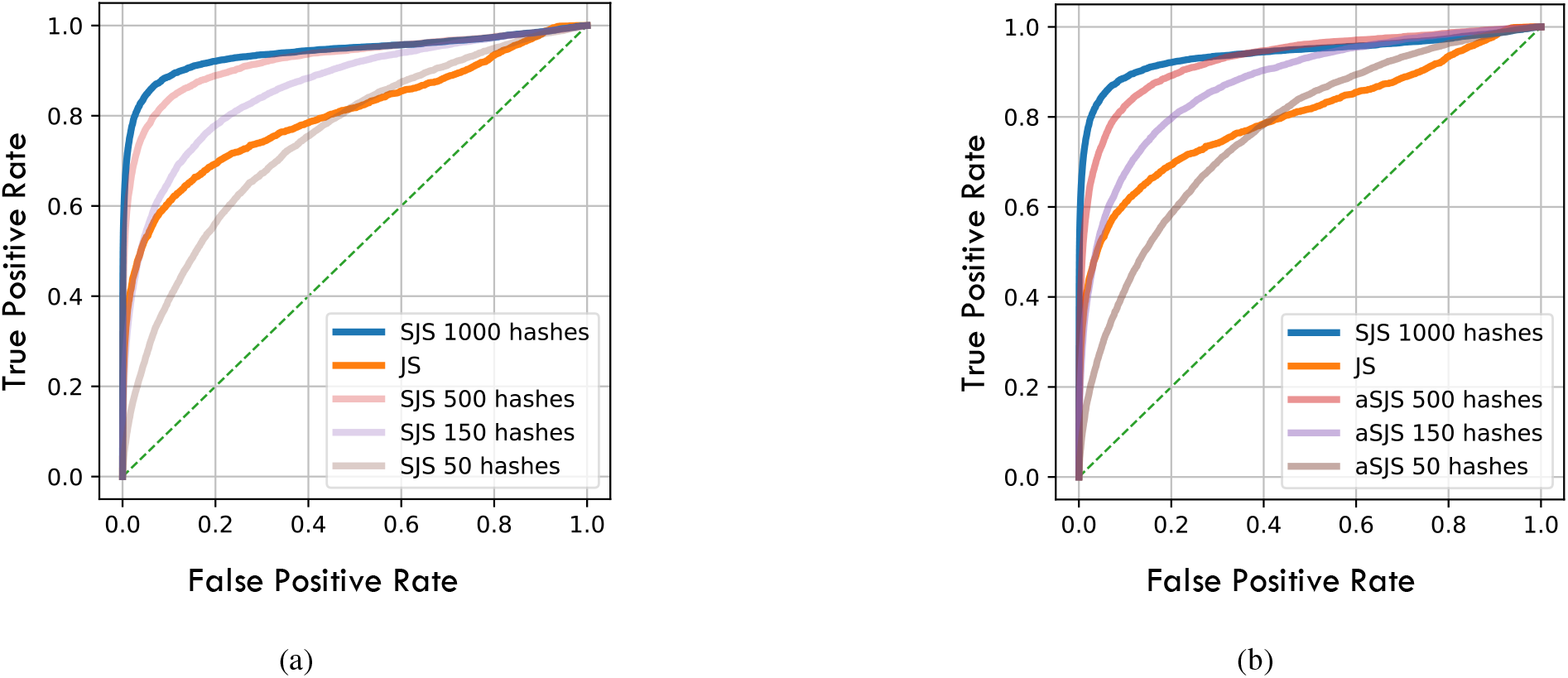
ROC curves obtained by (a) SJS and (b) aSJS as we vary the number of hash functions used for *θ* = 0.3 on Ecoli.

**Figure S18:**
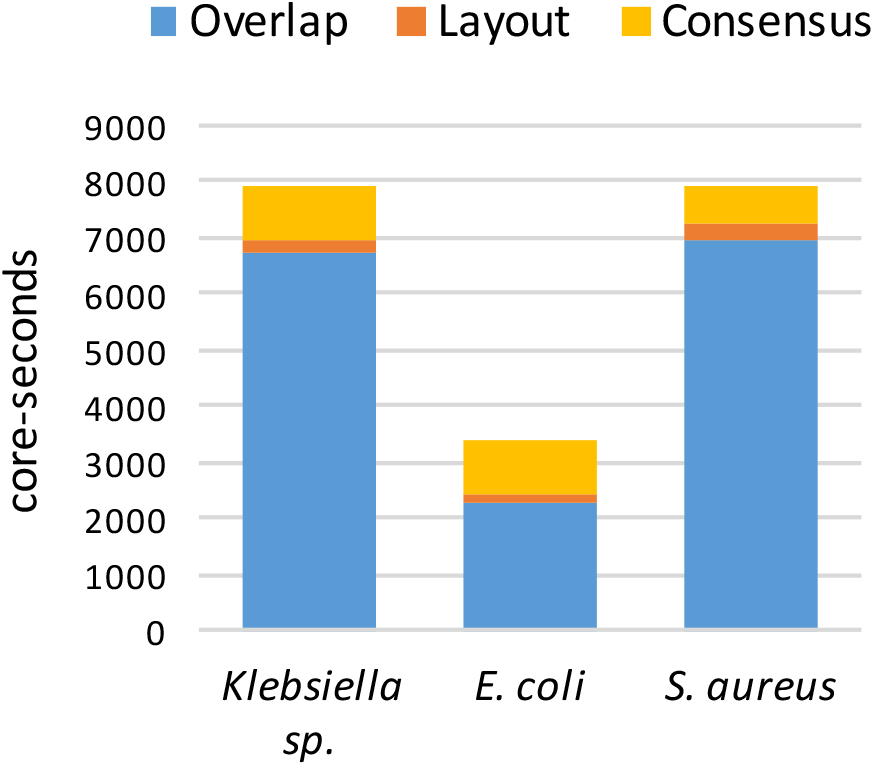
Time required for different stages of the assembly pipeline for different bacterial genomes. Note that the vast majority of time is spent determining pairwise alignments in the overlap stage (Adapted from Figure S11 in Kamath et al. [2017]).

## Notes

https://github.com/TavorB/spectral_jaccard_similarity

